# Using heterogeneity in the population structure of U.S. swine farms to compare transmission models for porcine epidemic diarrhoea

**DOI:** 10.1101/017178

**Authors:** Eamon B. O’Dea, Harry Snelson, Shweta Bansal

## Abstract

In 2013, U.S. swine producers were confronted with the disruptive emergence of porcine epidemic diarrhoea (PED). Movement of animals among farms is hypothesised to have played a role in the spread of PED among farms. Via this or other mechanisms, the rate of spread may also depend on the geographic density of farms and climate. To evaluate such effects on a large scale, we analyse state-level counts of outbreaks with variables describing the distribution of farm sizes and types, aggregate flows of animals among farms, and an index of climate. Our first main finding is that it is possible for a correlation analysis to be sensitive to transmission model parameters. This finding is based on a global sensitivity analysis of correlations on simulated data that included a biased and noisy observation model based on the available PED data. Our second main finding is that flows are significantly associated with the reports of PED outbreaks. This finding is based on correlations of pairwise relationships and regression modeling of total and weekly outbreak counts. These findings illustrate how variation in population structure may be employed along with observational data to improve understanding of disease spread.

## Introduction

The 2013 emergence of porcine epidemic diarrhoea (PED)^1^ in the United States has provided an example of both the economic hardships livestock diseases can cause and our limited understanding of how such diseases spread. Porcine epidemic diarrhoea virus (PEDV), the causative agent, acutely infects the intestine and causes severe diarrhoea and vomiting.^2^ Currently, the earliest known U.S. outbreak occurred in April,^3^ and in less than a year PED outbreaks were confirmed in 27 states,^4^ states that together produce 95 percent of the U.S. pig crop.^5^ Farms experiencing outbreaks have suffered 90 percent and higher losses of unweaned pigs.^3^ The time it takes for a farm to return to stable production is highly variable but on the order of weeks, leading to great expenses in infection control costs and production losses alike.

Losses were also apparent on a national economic scale. Producers had for the previous 8 years been making steady increases in the average litter size of about 0.16 head per year.^6^ By November 2013, the average litter size had begun an abnormal downturn,^6^ dropping 0.66 head by March 2014.^7^ The virus also affected swine production in Asia and other parts of America.^8, 9^

The mechanisms by which PEDV spread among farms are not yet clear. Transportation-associated transmission of PEDV has been supported by the observation at harvest facilities that it spreads among trailers used to transport swine,^10^ and some experts believe that current resources of livestock trailers, trailer-washing facilities, and transport personnel are insufficient to allow for a standard 3-hour trailer cleaning between every load.^11^ With such concerns in mind, some states responded to PED by requiring that imported swine be from PEDV-free premises. Transportation-independent mechanisms such as airborne particles^12^ and contaminated feed^13–16^ have also been implicated. Detailed investigations of outbreaks on farms can be inconclusive regarding the mechanism of PEDV introduction.^3^

Much of the research on PED involves detailed investigations on a small scale. For example, there have been epidemiological investigations of infected farms in North Carolina and a cluster of infected farms in Oklahoma and adjacent states.^17^ Such work is effective for determining the biological plausibility of different routes, but the risk factors identified in a small-scale study may be specific to the small area of the study. Modelling studies based on large-scale surveillance data^18–20^ can thus be a valuable complement to such work by quantifying the overall importance of a transmission route across a large population. Such quantification for PED could also be considered a contribution to the general study of infectious diseases of livestock. Although animal movements in general are considered a risk for transmission,^21^ only a limited number of studies^18–20,22,23^ have quantitatively compared this risk to other competing risks. One likely reason for this scarcity is that statistical analysis of the available data often presents many challenges such as incomplete and noisy reporting as well as correlations in explanatory variables.

Here we first evaluate the sensitivity of a correlation analysis to transmission and contact parameters of a simulation model of PED spread among U.S. swine farms via spatial and transport-associated pathways. The model includes an error-prone observational component designed to mimic that of the real data. We then apply this correlation analyis to the real data. We follow up on this analyis by considering a larger group of explanatory variables and applying stability selection to identify those with the most robust association with PED burdens. Using the selected variables, we formulate a simple model of farm-to-farm spread and estimate transmission parameters.

## Methods

### Natural history of PED outbreaks

We first provide a brief background on the natural history PED outbreaks to make clear the features that our models replicate and those that they do not. The time from the introduction of infected animals to the appearance of clinical symptoms in PEDV-naive herds is typically less than 7 days.^9, 24^ PED may spread rapidly within farms following the first appearance of clinical signs.^9,16,25–27^ The spread may also be actively promoted as a part of recommended feedback procedures to establish herd immunity.^24^ The duration of an outbreak can vary substantially. The lower bound would be roughly a week, as virus shedding from individuals has been observed in experimental settings to subside within 9 days of infection,^28^ and an infected animal’s diarrhoea has been observed in the field to typically last for 5 days.^25^ However, it can take affected farms several weeks to return to baseline production levels.^9,25^ In summary, within a farm spread is often rapid and complete enough that assuming all animals on the farm to have the same status is justified when observation occurs at weekly intervals.

### Structure of the U.S. swine herd

When we step back from a single farm and consider all the farms in the United States, a number of heterogeneities enter the picture. All of the contiguous 48 states share some portion of the nation’s swine but the Midwest and North Carolina are areas of major concentration (Supplementary Fig. S1), holding some 88 percent of the inventory.^5^ Thus considerable climactic and density gradients exist. Live swine are moved among farms for feeding and breeding purposes. We refer to aggregated measurements of these movements as transport flows. The value of these flows varies greatly between different pairs of states with the largest flows being into the Midwest (Supplementary Fig. S1). These heterogeneities may result in the spread of a disease resulting in a spatio-temporal distribution of outbreaks that is dependent on the mode of farm-to-farm spread, and for PED data on both the spatio-temporal pattern and the heterogeneities are available at the state level.

### Data sources

The data we analysed were obtained from public sources. The data on numbers of farms, balance sheets, and climatic regions came from regular USDA reports. The data on transport flows came from a study by the USDA Economic Research Service.^29^ The PED outbreak data were derived from a report published on the website of the American Association of Swine Veterinarians. A detailed description of these data may be found in the Supplementary Note. We next discuss an important assumption made about the PED outbreak data here.

Our working assumption is that the number of PEDV-positive accessions reported in 2013 is correlated with unknown number PEDV-positive farms. This assumption is necessary because because data on the number of PEDV-positive farms did not become available until June 2014. These more recent data do support our assumption that positive accessions and positive farms were correlated in 2013: positive accessions and positive farms have a Spearman rank correlation of 0.74 with data from June 2014 to February 2015.^30^ A plot of the counts of accessions and positive premises appears in Supplementary Fig. S2. Although it might seem preferable to analyse the 2014–2015 data on positive farms instead of positive accessions, the 2013 data may be more informative of transmission routes because farms protected by immunity rather than lack of exposure were most likely less frequent in 2013. Figure 1 shows the spatio-temporal pattern in the positive accessions.

**Figure 1.**
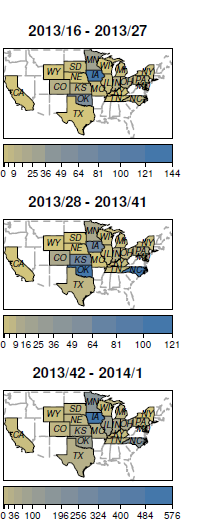
PEDV-positive accessions by state for three periods in 2013. Above each panel is the range of ISO weeks over which counts of positive accessions are aggregated to determine the fill color. The periods correspond roughly to spring, summer and autumn of 2013. Note that the scale of the colorbar is different for each panel and a lull occurred in the summer period for most states. States with no positive accessions in 2013 have dashed borders. This figure was created with the R package surveillance.^45^.

### Sensitivity analysis of simulated correlations

Since the data were highly aggregated and noisy, it was instructive for us to evaluate the performance of our correlation analysis under these conditions. To that end, we simulated the spread of a disease among individual U.S. swine farms, simulated a series of positive accessions, and then performed the correlation analysis on the simulated accession data. The model of disease spread included contact networks based on the spatial and transport structure of U.S. farms. The correlation analysis estimated the strength of association between the similarity of the time series of accessions between a pair of states with the measure of similarity according to either spatial or transport structure. Using previously described methods,^31,32^ we computed global sensitivity indices up to second order for the mean value of these correlations for the parameter space given in Table 1. These indices estimate the fraction of the variance in the correlation that is due to variation in individual parameters or pairs of parameters. This exercise did not validate our modeling assumptions, but rather evaluated self-consistency by showing the extent to which we could quantify and distinguish between different modes of spread given our assumed models of spread and observation. The procedure is described in further detail in the Supplementary Note.

**Table 1.** Space of input variables for the global sensitivity analysis.

| Variable name | Values | Description |
| --- | --- | --- |
| Spatial resolution | {1425, 570} | Columns in spatial grid (roughly 3 and 8 km grid cell sides) |
| Seasonal amplitude | 0–1 | Parameter controlling annual variation in transmission probabilities |
| Starting grid $x$ | 0–1 | East–West coordinate of region with first outbreak |
| Starting grid $y$ | 0–1 | North–South coordinate of region with first outbreak |
| $P(\text{trans. transport edge})$ | 0–1 | Probability of transmission along transport edge |
| $P(\text{trans. spatial edge})$ | 0–1 | Probability of transmission along spatial edge |
| $P(\text{external trans.})$ | $0\text{--}10^{-4}$ | Probability of transmission from external source |

### Stability selection

To see if important variables were missing from our correlation analysis, we performed variable selection for a regression model of the cumulative number of outbreaks in each state. Candidate predictors were chosen based on availability and expected effects on either reporting rates or risk. In addition to transport-associated risk, we considered that the risk of an outbreak may depend on climate, the density of farms in an area, and on a farm’s distance to other outbreaks. Table 2 contains a brief description of all the predictors considered and the Supplementary Note contains a detailed description.

**Table 2.** Variables available for selection in regression model of cumulative positive accessions and percent decrease in litter rates. This table summarises the variables used by describing groups of one or more variables that were closely related.

| Variable group | Description |
| --- | --- |
| Number of operations | Count of livestock operations in state with 25 or more swine. |
| Balance sheet | Dec. 2012 swine inventory and 2011–2012 pig crop, inshipments, and marketings. |
| Farm resource region | Proportion of swine farms in each state in each region, indicative of climate. |
| Nearby positive accessions | Weighted average of positive accessions nearby in flow network or geographically. |
| Farm density | Summary statistics for each state of the number of farms in each county per km<sup>2</sup>. |

Most of our predictors were correlated with other predictors as well as with the total positive accessions in each state. For such data, fitting regression models with an elastic net penalty allows groups of correlated variables to be given similar effect sizes whereas other modelling approaches, such as stepwise approaches and the use of a lasso penalty, may lead to one variable in a correlated group being singled out and being given a too-large effect size.^33^ In general for elastic net regression, the weight given to the penalty determines whether any variable is selected. Often, the goal of a regression analysis is to obtain a model with good predictive performance and the weight is chosen by cross validation.^33^ By contrast, we have no need of a predictive model and are instead more interested in determining what variables are important to include in a model. Stability selection^34^ provides a general method of identifying relevant variables. The main idea is to select variables that across many random subsamples of the data are selected with high probability by the elastic net with a given set of weights for the penalty. We use this procedure because it is less likely to select noise variables than is cross validation.^34^ Further details are given in the Supplementary Note.

### Time series regression modeling

To estimate how farm density and transport flows may affect transmission rates, we fit the parameters *c_i_* and *η* in the following regression model

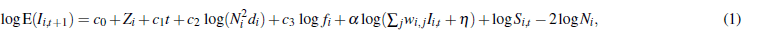

where E(*I*_*i, t*+1_) is the expected number of positive accessions in state *i* at week *t* + 1, *c*_0_ determines the baseline transmission rate, *Z_i_* is a normally distributed random effect on the transmission rate, *c*_1_ estimates the effect of seasonality on the transmission rate, *c*_2_ estimates the effect of farm density on the transmission rate, *N_i_* is the number of farms in state *i*, *d_i_* is the density of farms in state *i*, *α* captures nonlinearity created by clustering of infected farms in the contact network and weekly aggregation of the counts of a continuous process, *w_i, j_* are weights based on a submodel of how transport flows affect transmission rates, *I_i, t_* is the observed number of outbreaks in state *i* at week *t*, *η* is the rate of transmission from unobserved sources, and *S_i,t_* is the number of susceptible farms in state *i* at week *t*. For the submodels of how transport flows affect transmission rates, we considered an internal model assuming flows only affect transmission within a state, a directed model assuming that flows increased transmission from source states to destination states, and an undirected model assuming that transmission between a pair of states depended on the total flow in both directions. The derivation of these models as well as equation 1 is in the Supplementary Note.

We fitted these models to data from all 48 contiguous states with the assumption that the observed positive accessions *I*_*i,t*+1_ have a negative binomial distribution with an unknown, but constant, dispersion parameter which we denote with *θ*. This parameter is related to the variance by Var(*I*_*i, t*+1_) = E(*I*_*i,t*+1_)[1 + E(*I*_*i, t*+1_)/0]. We assumed that the random effect *Z_i_* is normally distributed. Then the likelihood is fully specified. We calculated marginal likelihoods with the Laplace approximation and numerically found the parameters that maximised it. In some cases we fixed *η* to 0.5, which allowed the model to be fully fit with both the lme4^35^ and glmmADMB^36^ packages in R.^37^ To make sure our results were not sensitive to *η* = 0.5, we used R’s optimise function to find the value of *η* in [0,5] with highest likelihood. We performed several diagnostic checks of our fits, including checking for signs of nonlinearity with partial residual plots and for signs of temporal autocorrelation in the residuals. We also verified that the flows term is significant in models lacking random effects, and after excluding any data points with dfbetas^38^ above 0.2.

## Results

### Pairwise correlations are sensitive to transmission and contact parameters

Before we present the correlation analysis on the empirical PED data, we first consider the sensitivity of the correlations to the parameters of a model of disease spread among farms in the U.S. swine herd. A large portion of the variance in our simulation output was stochastic. The value of 1 – *R*^2^ for our mean metamodel was 0.53, 0.46, and 0.18 respectively for correlations between the lag-1 cross correlation matrix and the distance, shared border, and transport flow matrices. Figure 2 shows that parameters with the highest sensitivities were the transmission probability for transport edges, the amplitude of seasonal variation in transmission probabilities, and the resolution of the raster grid used to generate spatial edges. All correlation types typically increased with transmission probabilities across transport edges, with those of the transport flow matrix increasing the most (Supplementary Fig. S3). All correlation types typically increased as spatial resolution decreased and the spatial contact network became more dense (Supplementary Fig. S4). Sensitivity to the spatial transmission probability was noticeable only on the more dense spatial network (Supplementary Fig. S5). Seasonal variation in the transmission probabilities tended to reduce correlations (Supplementary Figs. S3 and S4). Correlation size was indicative of statistical significance (Supplementary Figs. S3–S5). In summary, the largest sensitivities were to transmission and contact parameters, and these parameters had unsurprising relationships with the matrix correlations.

**Figure 2.**
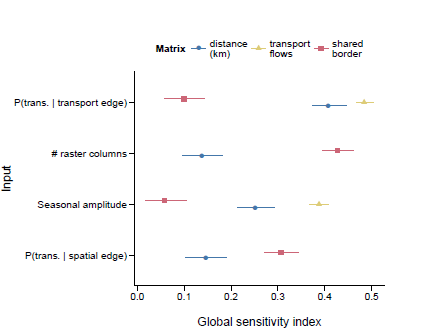
Matrix correlations were sensitive to disease model parameters. The response variable for the sensitivity analysis was the metamodel for the mean of the Spearman correlation between the matrix of lag-1 cross correlations of reported outbreaks and the matrix of state-to-state relationships indicated in the figure legend. Error bars are 95% confidence intervals. Sensitivity indices with confidence intervals that included zero were excluded from the plot.

### Similarity in outbreak count time series is correlated with transport flows

To provide insight into the relative importance of spatial and transport-associated transmission for PED, we applied the correlation analysis to data from the 2013 epizootic. We found that lag-1 cross correlations of positive accessions were positively correlated with the logarithm of transport flows. This relation held whether flows and cross correlations were treated as directional (Fig. 3a), were averaged over both directions (Supplementary Fig. S6), or were ranked (Supplementary Figs. S7 and S8). The size of the correlation was comparable to those seen in our simulations for high transmission probabilities across network edges (Supplementary Fig. S3). The *p* values for these correlations were all below 0.05 / 6, which means that they would remain significant after using a Bonferroni method of limiting the probability that any false positives occurred in our tests to 0.05. On the other hand, in no case would the correlations between geographic distance and cross correlations remain significant after such a correction. The correlations between the cross correlation matrix and the shared border matrix were similar to or weaker than those of the distance matrix, were not significant after Bonferroni correction, and were not included in the plots to keep them simpler. Analogous results hold for lag-0 and lag-2 cross correlations. Although higher lags are potentially informative, we did not attempt to analyse them as they were typically smaller, had more skewed distributions, and had greater uncertainty.

**Figure 3.**
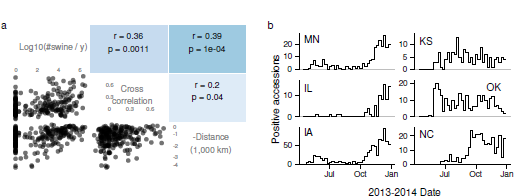
Similarity between state’s PED dynamics correlated more strongly with transport flows than with distance. (a) Scatter plots and Pearson correlations between transport flows, cross correlations between time series of positive accessions, and negative geographic distances. The *p* values are from a Mantel test. (b) PED dynamics for selected states. Distinct shapes are apparent in the time series of the Midwestern states (MN, IL, IA), Kansas and Oklahoma, and North Carolina.

The flows were themselves correlated with the distance between state centres, and these distances were in turn correlated with cross correlations (Fig. 3a). Thus we also calculated the partial correlation of flows and cross correlations, controlling for distance. This partial correlation was around 0.31 whether directed or undirected relationships were used, and thus controlling for distance does not greatly diminish the correlation.

### Numbers of farms and balance sheet variables were stable predictors of total outbreak counts

The Mantel test found a significant correlation between transport flows and cross correlations but did not account for many potential confounding variables. To address that limitation, we performed variable selection on a panel of candidate variables to identify those with the most robust associations with cumulative burdens of PED. Using stability selection with data from 42 observations, we found that among all variables in Table 2 the number of farms in a state was the only variable selected as a predictor of whether it reported any positive accessions. Among the 22 states reporting positive accessions, swine inventory and marketings were selected as predictors of the total number of positive accessions. Marketings is the total number of swine shipped out of a state or slaughtered.

### Transport flows and farm density are jointly associated with increased transmission

To obtain a rough estimate of the effect of flows on transmission rates that accounted for other effects such as seasonality and farm density, we fitted the case data to time series susceptible-infected-recovered models. Figure 4 displays the predicted and observed marginal relationships between flows and positive accessions for one of the models. Although our flow variables were based on the outcome of variable selection, they are not equivalent to any of the variables in the variable selection procedure and the data analysed here has a time dimension not present in the data used for variable selection. Thus to confirm the statistical significance of the within-state flows, we conducted a likelihood ratio test of the hypothesis that models lacking terms for within-state flow were sufficient. The test favoured rejection of models without within-state flows 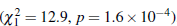.

**Figure 4.**
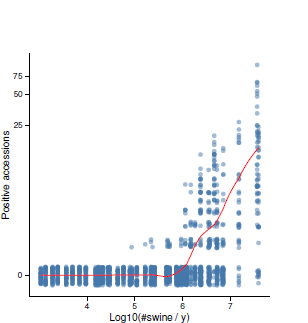
Transport flows were predictive of the number of new positive accessions. The line is a LOESS smoother of predicted values from the undirected model, where predictions were calculated for each observation by adding fixed effects and conditional modes of random effects. The points are the original data. To display their density, they have been made transparent and jittered along the y axis. The *y* axis was transformed using *y* = log(Positive accessions+1).

Among those models containing flows, undirected models, which assumed that flows increased contact rates in both source and destination states, fit best, and directed models, which assumed that flows increased contact of susceptible farms in the destination state to infective farms in the origin state, fit worst (Table 3). However, the parameter estimates were generally similar for all of these models, with flows having an appreciable effect (Fig. 5).

**Table 3.** Summary of models. The models chiefly differ by how contact is assumed to depend on flows. In the null model, denoted by none, contact was independent of flows. In the internal model, contact was a function of within-state flows. In the directed model, contact was a function of flows moving into a state and within-state flows. In the undirected model, contact was a function of within-state flows and both flows into and out of a state. The column “Fit *η*?” indicates whether we estimated the value of *η*, which corresponds to risk that is independent of the number of infective farms. The symbol *θ* denotes the dispersion parameter of the negative binomial response. The symbol *σ* denotes the standard deviation of the random effect of (geographic) state on transmission rates. The abbreviation d.f. is for degrees of freedom (i.e., the number of parameters estimated). ΔAIC gives the AIC (Akaike information criteria) of a model minus the lowest AIC of all models.

| Flow term | Fit $\eta$? | Intercept | $\hat{\theta}$ | $\hat{\sigma}$ | d.f. | Log lik. | $\Delta\text{AIC}$ |
| --- | --- | --- | --- | --- | --- | --- | --- |
| undirected | yes | −4.5 | 2.21 | 1.36 | 8 | −999.8 | 0.0 |
| directed | yes | −4.7 | 1.94 | 1.52 | 8 | −1017.9 | 36.3 |
| internal | yes | −4.0 | 2.20 | 1.42 | 8 | −1005.4 | 11.3 |
| internal | no | −3.8 | 2.17 | 1.40 | 7 | −1005.6 | 9.7 |
| none | no | −3.9 | 2.16 | 1.82 | 6 | −1012.1 | 20.6 |

**Figure 5.**
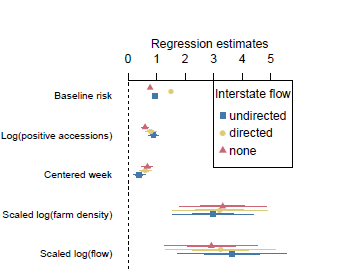
Parameter estimates for the transmission model. The estimates are not sensitive to the choice of interstate flow model, and flows have similar effect sizes to farm densities. Baseline risk refers to the parameter *η*, which determines the risk of infection when no infectives are present. The error bars represent 50- and 95-percent Wald confidence intervals. The scaled variables were divided by the interquartile ranges to make their effect estimates comparable. The interquartile ranges for week and the logarithm of farm density were 19.0 and 4.5. Those for the logarithm of undirected, directed, and internal (no interstate) transport flows were 4.5, 4.3, and 3.8.

## Discussion

Our first main finding is that our correlation analysis method may for a range of parameter values remain sensitive to transmission and contact parameters of interest in spite of receiving partial, biased, and noisy data about the state of simulated epizootics as input. In short, the signals generated by the heterogeneity in the structure of the swine herd were not drowned out by the substantial noise generated by the observation model. The potential of herd structure to strongly influence the dynamics of swine diseases has been noted in previous work,^39^ and our results illustrate how from the dynamics it may be possible to identify the elements of herd structure relevant to the spread of a disease. The extent to which this identification is practical could be further clarified by developing a model including more realistic farm distributions,^40^ transportation networks, and observation models.

Our second main finding is that variables related to transportation network of the U.S. swine herd appear relevant to the dynamics of PED based on correlation analysis, stable associations with cumulative burdens, and a time series regression model. The idea that transportation is associated with the risk of PED transmission is not new, but our analysis does provide a new argument in support of it as well as parameters for a model of spread via transportation based on field data. From Fig. 5 we have the estimate for the directed model that one factor in the average pairwise transmission rate from farms in one area to those in another increases with the annual transport flow raised to the power of about 3.2/4.3 ≈ 0.75. In general, transmission rate parameters have a strong effect on the output of models of livestock disease spread and modellers must rely on expert opinion to set them.^41,42^ Estimates such as ours may thus be key for determining what parameter values are consistent with past epizootics.

What features of the time series might have driven the results in our correlation analysis? The correlation between transport and cross correlations seemed to be driven in part by concentration of both high cross correlations and large flows in Midwestern states (Supplementary Fig. S1). The cross correlations of these states results from the presence of a small wave of positive accessions early in the outbreak and a much larger wave toward the end of our observations (Fig. 3b, left column). Also, Kansas and Oklahoma share a distinctive period of high positive accessions in the middle of the time series and fairly large flows (Supplementary Fig. S9 and Fig. 3b). Epidemiological reports suggest that windborne transmission was important for spread in Oklahoma, and thus the summer activity in Kansas and Oklahoma may also have been a consequence of short-distance, spatial spread. It is not known how the spread in these states occurred in spite of the high temperatures that were thought to have slowed PEDV transmission in other states during the summer. North Carolina is distinct from the Midwest in that no outbreaks occurred in the spring and many outbreaks occurred in October (Fig. 3b). Epidemiological reports suggest that PEDV was introduced in the Midwest and eventually reached North Carolina via transport. The high number of sow farms in the state may have allowed for a largely self-sustained cluster of outbreaks following introduction. Although the matrices measuring spatial transmission may not have come out as significant in our analysis, consideration of the epidemiological explanations for the time series suggests that spatial transmission may to some extent explain the significance of the transport flow matrix.

Both epidemiological and statistical mechanisms may explain why the undirected model fit best (Table 3) and why pair-averaged flows had higher correlations than directed flows (e.g., Fig. 3 and Supplementary Fig. S6). A possible epidemiological mechanism is that trucks arriving to pick up loads are introducing the virus to farms. A possible statistical mechanism was seen in our simulations. We noticed that even though transport contact was based on directed transport flows, symmetrised matrices had slightly higher correlations than asymmetric matrices.

Our analyses were likely limited in power by inaccuracies in our variables measuring population structure. For example, the transport flows used excluded transport to harvest plants, and such movements have been observed^10^ to result in the contamination of trailers. Another concern is the coarseness and age of the flow estimates. In support of them being sufficiently informative, previous phylogeographic analysis^43^ has found evidence that the same flow data we used was predictive of the movement of H1 influenza A virus among swine. This result suggests that in spite of ongoing change in the population structure of the swine herd the 2001 flow estimates have a relatively stable predictive value because the samples for that phylogenetic analysis came from years 2005–2010. Likewise, 2009 interstate flow estimates for the U.S. cattle herd were in general agreement with 2001 estimates.^44^ Of course, updated flow estimates are desirable for future modelling.

The main limitation of this analysis is that flows are correlated with several other variables, and we cannot rule out that these other variables are the true drivers of the observed effect of flows. We have formulated our time series SIR model based on the relationship between flows and inventory. These two variables are closely related both because more swine are moving through the farms with larger inventory and because swine often have shorter residence times on larger farms, since larger farms tend to specialise on specific production stages. But if larger farms did not experience more PED outbreaks but only reported them with higher probability, that could provide a false signal that flows are associated with risk. A phylogeographic analysis or analysis of suitably structured epidemiological data could establish an association between flows and spread of PEDV that is not subject to such confounding.

## Conclusions

Both the objectives, identifying variables relevant to the risk of infection, and challenges of our data analysis, uncertain reporting rates and many correlated candidate predictors, are common in epidemiological studies. Two reasonable steps toward such an objective are to assemble as many relevant explanatory variables as possible about reporting rates and measures of exposure based on prior scientific knowledge and then to determine if the available data support the conclusion that these variables are relevant. Our general contribution has been to provide a worked-out example of how variation in the structure of the population across a large scale may allow for the identification of variables with relevance to mechanisms of spread. We have also demonstrated the use of stability selection and regularised regression for the task of filtering out noise variables from a set of candidates. These examples may serve to provide analysts with new ideas about how to make the most efficient use of often limited epidemiological data, hopefully leading to more rapid understanding of transmission and how to stop it.

## Acknowledgments

This work was supported by the RAPIDD Program of the Science & Technology Directorate, Department of Homeland Security and the Fogarty International Center, National Institutes of Health; as well as the Department of Homeland Security Science & Technology Directorate Foreign Animal Disease Modeling Program through contract # HSHQDC-12-C-0014.

We thank John Korslund for useful feedback on veterinary subject matter. We thank John Drake and Chris Dibble for comments that improved the clarity of the writing.

## Author contributions statement

S.B. and E.O. designed the study and drafted the manuscript. E.O. conducted the analyses. H.S. reviewed the veterinary subject matter of the manuscript.

## Additional information

**Competing financial interests**: The authors declare no competing financial interests.

